# DNA Methylation Network Estimation with Sparse Latent Gaussian Graphical Model

**DOI:** 10.1101/367748

**Authors:** Bernard Ng, Sina Jafarzadeh, Daniel Cole, Anna Goldenberg, Sara Mostafavi

**Affiliations:** Centre for Molecular Medicine and Therapeutics, UBC, Canada; SickKids Hospital, University of Toronto, Canada

## Abstract

Inferring molecular interaction networks from genomics data is important for advancing our understanding of biological processes. Whereas considerable research effort has been placed on inferring such networks from gene expression data, network estimation from DNA methylation data has received very little attention due to the substantially higher dimensionality and complications with result interpretation for non-genic regions. To combat these challenges, we propose here an approach based on sparse latent Gaussian graphical model (SLGGM). The core idea is to perform network estimation on *q* latent variables as opposed to *d* CpG sites, with *q*<<*d*. To impose a correspondence between the latent variables and genes, we use the distance between CpG sites and transcription starting sites of the genes to generate a prior on the CpG sites’ latent class membership. We evaluate this approach on synthetic data, and show on real data that the gene network estimated from DNA methylation data significantly explains gene expression patterns in unseen datasets.

## 1 Introduction

Uncovering networks of interacting genes provides insights into the biological mechanisms that give rise to phenotypes. Under this network perspective, genes correspond to nodes with their interactions modeled via weighted edges. Over the past decade, gene interaction networks have primarily been constructed from gene expression data due to their large abundance [1]. In contrast, less effort has been placed on using DNA methylation data, despite their relatively more robust (less dynamic) nature and increasing availability for large cohorts. Most studies that do use methylation data estimate networks by directly correlating all CpG site pairs, with a focus on module detection [2–6]. However, the typical small sample-to-variable ratio limits the accuracy of the resulting networks [7]. Also, interpreting methylation networks is more difficult, since less is known about the functional role and gene targets of non-coding regulatory regions. Some studies employ canonical correlation analysis (CCA) to combine gene-proximal CpG sites in reducing dimensionality and enabling gene level interpretation [8; 9], but CCA cannot distinguish direct interactions from indirect interactions. To address the above challenges, we propose here an approach based on sparse latent Gaussian graphical model (SLGGM) [10]. The idea is to estimate a network between *q* latent variables as opposed to *d* CpG sites, and tie the latent variables to genes via a prior on the CpG-to-gene mapping. This way, the scale of the network estimation problem is greatly reduced and gene level interpretation is facilitated. Also, SLGGM inherently estimates partial correlation, which helps isolate direct interactions [11]. We assess this approach on synthetic data, and further show that the gene network constructed from a large-scale methylation dataset (ROSMAP [12]) significantly explains gene expression patterns of unseen datasets from various related tissues (GTEx [13]).

## 2 Methods

### 2.1 Sparse Latent Gaussian Graphical Model

Given a *n×d* DNA methylation data matrix, **X**, where *n* is the number of samples and *d* is the number of CpG sites, our goal is to learn a *q×q* sparse inverse covariance matrix, **K**, where *q<d* and **K**_*ij*_≠0 indicates that genes *i* and *j* are conditionally associated with each other, i.e. only direct interactions are captured. This problem can be formulated as finding a **K** that maximizes the following distribution [10]: 

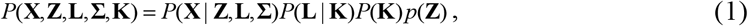

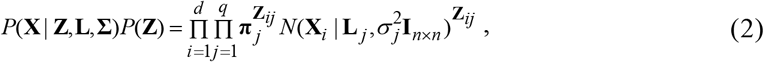

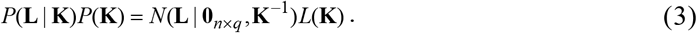

Each column of **X**, denoted as **X**_*i*_, is assumed to follow a mixture of Gaussians with means **L** = (**L**_1_,…,**L**_*q*_) and variances 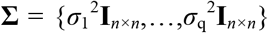. The *n*×*q* latent variable **L** is assumed to follow a centered multivariate Gaussian with an inverse covariance **K** having a Laplace prior *L*(**K**) to promote sparse **K** estimation, since *n<<q* typically. **Z** is a *d*×*q* binary matrix, where **Z**_*ij*_=1 indicates that **X**_*j*_ belongs to latent class *j*. Each **X**_*i*_ is assumed to belong to only one latent class as required for standard Gaussian mixture models. Optimal **L**, **∑**, and **K** can be found with coordinate ascent [10], i.e. alternate between finding **L**, **∑**, and **K** that maximizes (1). The update procedures for **L**, **∑**, and **K** are described in Sections 2.2, 2.3, and 2.4, respectively.

To deploy SLGGM for learning a gene network from methylation data, we impose a distance-based prior by setting ***γ**_ij_* = *E*(**Z**_*ij*_) to *f*(1/**D**_*ij*_), where **D**_*ij*_ is the base-pair distance between CpG site *i* and the transcription starting site (TSS) of gene *j*, and *f*(·) normalizes 1/**D**_*ij*_ such that ∑_*j*_ ***γ**_ij_* =1. The assumption is that CpG sites closer to a given gene are more likely to affect that gene, as often observed [14]. In this work, we fix ***γ**_ij_* to retain the correspondence between the genes and the latent variable L, and leave updating ***γ**_ij_* to refine the CpG-to-gene mapping for future work. Also, we note that in contrast to methylation values per sample, which tend to be bimodal, the distribution of methylation values at each CpG site is by and large unimodal and approximately Gaussian in the real data, hence meeting the assumptions of SLGGM.

### 2.2 L Update

By taking the log of (1), retaining terms involving **L**, and differentiating w.r.t. **L**_*j*_, one can show that the optimal **L**_*j*_ at iteration *k*+1 is given by: 

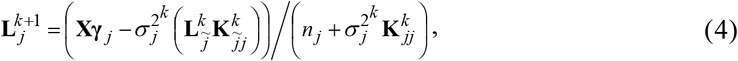

where ***γ*** = (***γ***_1_,…,***γ**_q_*) is a *d*×*q* matrix containing the distance-based prior, *n_j_* = ∑_*i*_ ***γ**_ij_*, and 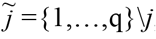, i.e. 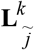 denotes all columns of **L**^*k*^ except the *j*^th^ and 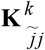 denotes the **j**^th^ column of **K**^*k*^ excluding the *j*^th^ element. We initialize **L** as **X*γ***.

### 2.3 *σ_j_*^2^ Update

Similarly, differentiating the log of (1) w.r.t. *σ_j_*^2^, the optimal *σ_j_*^2^ is given by: 

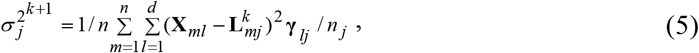

where the noise variance of **X** is assumed to be the same across the *n* samples for each latent class *j*. *σ_j_*^2^ is initialized to 1.

### 2.4 K update

Finding **K** that maximizes (1) is equivalent to solving the following problem [10]: 

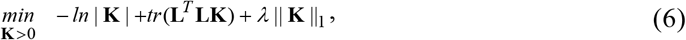

where the first two terms correspond to −log(*N*(**L**|0_*n×q*_,**K**_-1_)) with constants removed, and the last term is the *l_1_* norm of **K**_*ij*_, *i*≠*j*, corresponding to −log(*L*(**K**)), which promotes a sparse **K** estimate. For solving (6), we employ QUIC [15] and BigQUIC [16], depending on *q. λ* is set to *αλ*_max_ for different *α*’s, where *λ*_max_=*max_ij_* |**L**^T^**L**|, *i*≠*j*.

## 3 Materials

### Synthetic data

We created 60 synthetic datasets for three *q*/*n* ratios: 1=100/100, 5=500/100, and 0.5=500/1000, with 20 datasets for each *q/n* ratio. For each dataset, we first generated a random *q*×*q* sparse inverse covariance matrix, **K**, with 10% of the elements being non-zero. We then drew a random *n*×*q* matrix, **L**, from *N*(**0**_*n*×*q*_, **K**^-1^). A *n*×1 vector, **X**_*j*_, was then drawn from *N*(**L**_*j*_,*σ_j_*^2^**I**_*n*×*n*_) for *i*=1 to *d*, where the choice of *j* was based on **Z**_*ij*_=1. The CpG-to-gene mapping, **Z**, was established by assigning each CpG site to its closest gene based on the real data. On average, *d*≈3,000 and 15,000 CpG sites for *q*=100 and 500 random genes. The membership prior, ***γ**_ij_*, was set to *f*(1/**D**_*ij*_), where **D**_*ij*_ is the distance between CpG site *i* and TSS of gene *j*, and *f*(·) ensures ∑_*j*_ ***γ**_ij_*=1. Since **Z** is supposedly unknown, we included **D**_*ij*_ of all CpG-gene pairs that are within 1Mb from each other in computing ***γ*** to emulate uncertainty in the CpG-to-gene mapping. These smaller scale problems permit rigorous testing of SLGGM within a practical amount of computation time.

### Real data

DNA methylation data (450K Illumina array) were generated from 702 post-mortem samples of the dorsolateral prefrontal cortex as part of the ROSMAP study [12] (available on Synapse). In addition to standard quality control, age at death, sex, batch, post mortem interval, and top 10 principal components (PC) from the DNA methylation data were regressed out. For method evaluation, we used gene expression data (13,484 genes) from 508 individuals of the ROSMAP study [17]. Standard quality control was performed, and age at death, sex, batch, post-mortem interval, RNA integrity, sequencing depth, genotyping PCs, and top 10 PCs from expression data were removed. Only genes and CpG sites within 1Mb from each other were retained, resulting in 416,452 CpG sites and 13,004 genes. We also downloaded gene networks constructed from expression data of 13 brain tissues as well as expression data of 35 peripheral tissues from the GTEx portal [13].

## 4 Results and Discussion

### Synthetic data

Average precision and recall estimated by applying SLGGM (with QUIC), and comparing the sparse edge patterns of the ground truth and estimated networks are plotted in Fig. 1. For comparison, we applied SGGM (with QUIC) to the ground truth **L**. The performance of SLGGM applied to **X** is similar to SGGM applied to **L** for the *q*/*n* ratios tested, with SLGGM’s performance being well within the standard error of SGGM (not plotted to avoid clutter). If we weight the ground truth edges by |**K**_*ij*_| in the precision and recall estimation, both SLGGM and SGGM achieved a precision of ∼1, indicating that the stronger edges were reliably extracted.

**Fig. 1:**
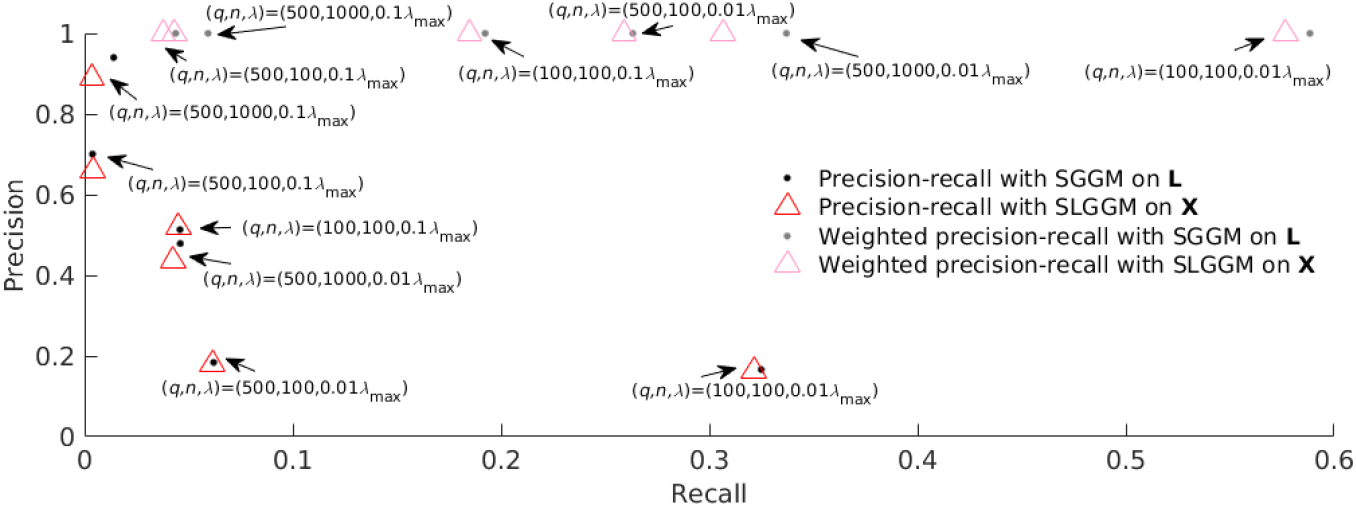
Synthetic data results. For various *q*/*n* ratios, applying SLGGM on **X** (*n*×*d*) resulted in similar performance as applying SGGM on the ground truth **L**(*n*×*q*), where *d*>>*q*.

### Real data

Since no ground truth network is available for real data, we used gene networks estimated from ROSMAP and GTEx expression data for assessing SLGGM’s performance. With the ROSMAP gene expression data, we applied SGGM (BigQUIC, *λ*=0.5*λ*_max_) and stability selection [18] on 100 random subsamples (0.8*n* subjects each). Network edges with selection frequency >0.5, i.e. estimated to be non-zero for >50% of the subsamples, were assumed reliable. With the 13 GTEx brain tissue-based networks, we assumed edges that were non-zero for >50% of the networks were reliable. As a baseline, we compared the reliable network edges between ROSMAP and GTEx. With ROSMAP as the reference, the estimated precision and recall are 0.023 and 0.061. If we weight the reliable network edges by the selection frequency, the weighted precision and recall are 0.917 and 0.062, demonstrating that the more reliable edges tend to be extracted. Comparing the network edges extracted by SLGGM (BigQUIC, *λ*=0.2*λ*_max_) and the reliable edges in the ROSMAP and GTEx expression networks, the precision and recall are 0.001 and 0.059 for ROSMAP and 0.001 and 0.023 for GTEx. The low precision is likely due to the small sample size. Nonetheless, the weighted precisions of SLGGM are 0.920 for ROSMAP and 0.306 for GTEx, indicating that the more reliable network edges are indeed extracted. We note that assessing only the sparse edge patterns neglect the information encoded by the edge weights. To capture this information, we applied kernel machines [19] with the ROSMAP and GTEx expression data as response and **K**^-1^ estimated with SLGGM as the kernel. Specifically, we averaged the expression values across subjects for each gene but without subject mean removal to retain the genome-wide expression pattern. This pattern of average expression values was used as the response. The association between the ROSMAP expression pattern and **K**^-1^ is statistically significant (*p*=0 to machine precision), demonstrating that **K**^-1^ well explains the gene expression pattern. Also, over the 35 tissues in the GTEx data, stronger associations are observed for brain tissues on average, Fig. 2.

**Fig. 2:**
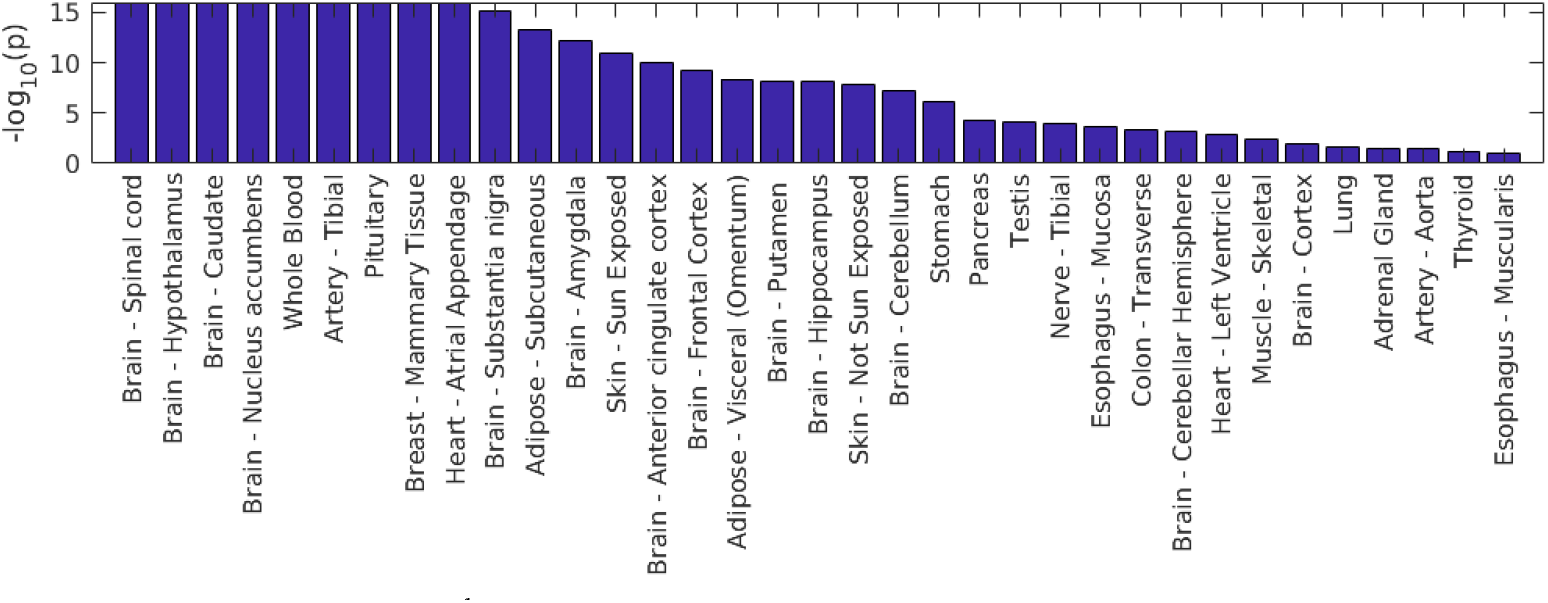
Real data results. The **K**^-1^ estimated by SLGGM well explains the expression pattern of brain tissues. Note p-values that are 0 (to machine precision) were set to 10^-16^ for clearer visualization.

